# Antiproliferative Effect of *Epilobium parviflorum* Extracts on Colorectal Cancer Cell Line HT-29

**DOI:** 10.1101/2020.12.12.422402

**Authors:** M. Aydın Akbudak, Tevhide Sut, Nuraniye Eruygur, Ersin Akinci

## Abstract

The *Epilobium* species are rich in various active phytochemicals and have seen wide use in folk medicine to treat several diseases. The present study exhibited the antiproliferative activity of aqueous and ethanolic *Epilobium parviflorum* extracts in a colon cancer cell line, HT-29 cells. Both types of extracts reduced the cell viability of HT-29 cells in a dose-dependent manner. A gene expression analysis of the HT-29 cells demonstrated an increase in apoptotic genes, *Caspase-3* and *Caspase-8*. Nuclear fragmentation of the apoptotic cells was also demonstrated through TUNEL assay and immunostaining experiments. On the other hand, the same lethal concentrations of the *E. parviflorum* extracts did not significantly affect a non-cancerous human fibroblast cell line, BJ cells. Our results confirmed that aqueous and ethanolic *E. parviflorum* extracts can eliminate proliferation of human colorectal carcinoma cells *in vitro*. *E. parviflorum* may have the potential to become a therapeutic agent against colon cancers.

## Introduction

The *Epilobium* genus from the *Onagraceae* family contains more than 200 species that are distributed all around the world. Among them, *E. angustifolium* L., *E. parviflorum* Schreb., and *E. hirsutum* L. are the most well-known (Vitalone and Allkanjari 2018) and are widely used in traditional medicine to treat various infections, disorders and diseases. To illustrate, their extracts have served as remedies to relieve menstrual disorders in Chinese folk medicine and to treat infected sores, swellings and rectal hemorrhages in traditional Native American medicine (Granica et al. 2014; Hevesi et al. 2009; Stolarczyk et al. 2013). *E. angustifolium*, *E. montanum* and *E. parviflorum* have also been used to cure prostate, kidney and urinary tract diseases in Austrian folk medicine (Vogl et al. 2013). Moreover, they have been reported to treat skin and mucosa infections (Granica et al. 2014; Stolarczyk et al. 2013). In parallel with their use in folk medicine, the aerial parts of the *Epilobium* species have been demonstrated to be rich in various active phytochemicals, such as phytosterols, polyphenols, flavonoids, and tannins. It was reported that *Epilobium* sp. contains 103 different compounds (reviewed by Granica et al. (2014)). According to ultra violet (UV) and tandem mass spectrometry (MS/MS) analyses, the phytochemical characterizations of *E. angustifolium*, *E. parviflorum*, and *E. hirsutum* extracts are comprised of 38 compounds (Stolarczyk et al. 2013).

In addition to the disorders and diseases mentioned above, extracts from the *Epilobium* species are most commonly used as infusions to treat benign prostatic hyperplasia (BPH), urethral inflammation, and micturition disorders. BPH is the most significant urological disorder that affects the aging male population because it causes excessive cellular proliferation in prostrate tissues (Vitalone et al. 2001). Although its etiology is not fully known, BPH is regarded as an endocrine disorder caused by an age-related hormone imbalance (Ekman 1989). Previous studies have demonstrated the inhibitory activity of *Epilobium* spp. extracts against 5α-reductase (which converts testosterone to dihydrotestosterone) and aromatase (which converts testosterone to estrogen) enzymes, which are involved in the etiology of BPH (Ducrey et al. 1997). Oenothein B, the primary tannin component of *E. parviflorum*, is proven to inhibit 5α-reductase enzymes. Although BPH can be cured through medical operations, diuretic problems are often reported due to insufficient resection and persistent urinary infections. Therefore, *Epilobium* herb infusions remain an attractive alternative for the treatment of BPH (Akbudak 2002).

In addition to the anti-androgenic effects of *Epilobium*, the anti-cancerous effects of the genus’ extracts have been demonstrated on various cell lines, including androgen-independent prostate cancer cells (PC-3), androgen-sensitive human prostate adenocarcinoma cells (LNCaP), human mammary epithelial cells (HMEC), the human astrocytoma cell line (1321 N1), and human epithelial prostrate cells (PZ-HPV-7) (Kiss et al. 2006; Stolarczyk et al. 2013; Vitalone et al. 2001; Vitalone et al. 2003b).

Colorectal cancer is one of the deadliest forms of cancer. While it can be cured by surgical removal of the tumor if detected early, effective treatment options become increasingly limited as the disease progresses. In addition to colon cancer’s links to genetic predisposition, aging, smoking and a lack of exercise, reports have indicated that it is also connected to diet and that at least some colon cancers can be prevented by changing dietary habits. In light of the pathogenesis of prostate cell proliferation and the application of *Epilobium* sp. extracts in folk medicine and their activities on various cancer cell lines, we investigated the antiproliferative activity of *E. parviflorum* extracts obtained from the solvents of water and ethanol on a human colorectal adenocarcinoma cell line (HT-29).

## Material and Methods

### Plant material and extract preparation

*E. parviflorum* seeds were purchased from The Strictly Medicinal Seeds Company (OR, USA). Seeds were individually sown into 20-cm square pots containing peat. Plants were grown under a regime of 16h-light/8h-dark periods at 24°C and 60% relative humidity in a plant growth chamber. They were watered with half-strength Hoagland’s liquid medium every day. After reaching maturity (two-months old), the aerial parts of the plants were harvested and shade dried at room temperature. Thirty-five grams of the powdered plant material was macerated with 50% ethanol for 24 hours. After filtering by filter paper, the extraction was repeated two times. The combined filtrate was concentrated under vacuum at 40 °C by rotary evaporator (Buchi R-100, Switzerland) to obtain the crude aqueous ethanol extract (13.61 g, Yield: 38.89%). A crude ethanolic extract was suspended in distilled water (500 mL) and was then fractioned with n-hexane (10 × 500 mL), ethyl acetate (10 × 500 mL), n-butanol (10 × 500 mL), and distilled water (10 × 500 mL) using separating funnels. All fractions were concentrated using a rotary evaporator, from which 0.23 g hexane frc. (2.21%), 2.09 g ethyl acetate frc. (20.1%), 2.16 g n-butanol frc. (20.77%), and 5.095 g aqueous frc. (48.99%) were obtained (Ayaz et al. 2014).

### Cell Culture

Human colon cancer cell line HT-29 and noncancerous human fibroblast cell line (BJ) were purchased from ATCC (VA, USA). The cells were plated into T-75 tissue culture treated flasks (Corning, NY, USA) with DMEM (Gibco, MA, USA) medium supplemented with 10% fetal bovine serum (FBS) (Biological Industries, VT, USA), 1% antibiotic-antimycotic (Gibco), 1% L-glutamine (Gibco) and maintained at 37 °C in a 5% CO_2_ humidified environment. The cells were passaged using trypsin (Gibco) once the plate became confluent. Before the plant extract application, the cells were seeded into a 96-well plate (Corning) as 10^4^ cells in each well and then incubated for 24 hours before the application. Powdered plant extracts of *E. parviflorum*, obtained using 2 different solvents (water and ethanol), were dissolved in distilled water. Ethanolic plant extract was diluted in the medium at a concentration of 500, 200, 100, 10 μg/ml. Aqueous plant extract was diluted to 100, 50, 30, 10 μg/ml in the medium. Dilutions were then applied onto HT-29 and BJ cells and incubated for 48 and 72 hours.

### MTT (Cell Proliferation Assay)

HT-29 and BJ cells were plated into 96-well plates at 10^4^ cells/well. The MTT (Millipore, MA, USA) solution was prepared at a concentration of 5 mg/ml. Following the application of plant extracts, 30 μl of stock MTT was added into each well on day 2 and day 3. Plates were incubated at 37 °C for 4 hours. After incubation, the medium was removed and 100 μl of DMSO was added into each well to dissolve the purple formazan. Using a spectrophotometer, formazan was read at 570 nm and the background was read at 690 nm. Based on the control group, the percentage of viable cells was calculated using the absorbance values found. The experiments were conducted with three biological and three technical replicates. The half maximal inhibitory concentration (IC_50_) values were calculated by GraphPad Prism 7 software.

### RNA Isolation and cDNA Synthesis

Total RNA isolation was performed using GeneJET RNA Purification Kit (Thermo Scientific, MA, USA) according to the manufacturer’s protocol. Any genomic DNA contamination was removed using RNase-Free DNase enzyme (Promega, WI, USA). RNA amounts were determined with BioDrop μLITE (Biodrop, UK). RNA samples were stored at −80 °C. 500 ng of total RNA from all samples were transcribed into cDNA using iScript cDNA Synthesis Kits (Bio-Rad, CA, USA) according to the manufacturer’s protocol. The resulting cDNA samples were stored at −20 °C.

### Quantitative PCR Assay

The level of gene expression was measured using LightCycler 96 system (Roche, Germany) and Maxima SYBR Green RT-qPCR Master Mix (Thermo Scientific). Sequences of *Caspase-3*, *Caspase-8*, *p53*, *Bax*, and *glyceraldehyde-3-phosphate dehydrogenase (Gapdh)*, genes were obtained from NCBI database and the primers (Table 1) were designed using NCBI primer software. The specificity of the primers was confirmed by a single band image on the agarose gel. *Gapdh* gene was used as the reference. Gene expression levels were demonstrated using ΔΔCT method (Livak and Schmittgen 2001).

**Table 1.**
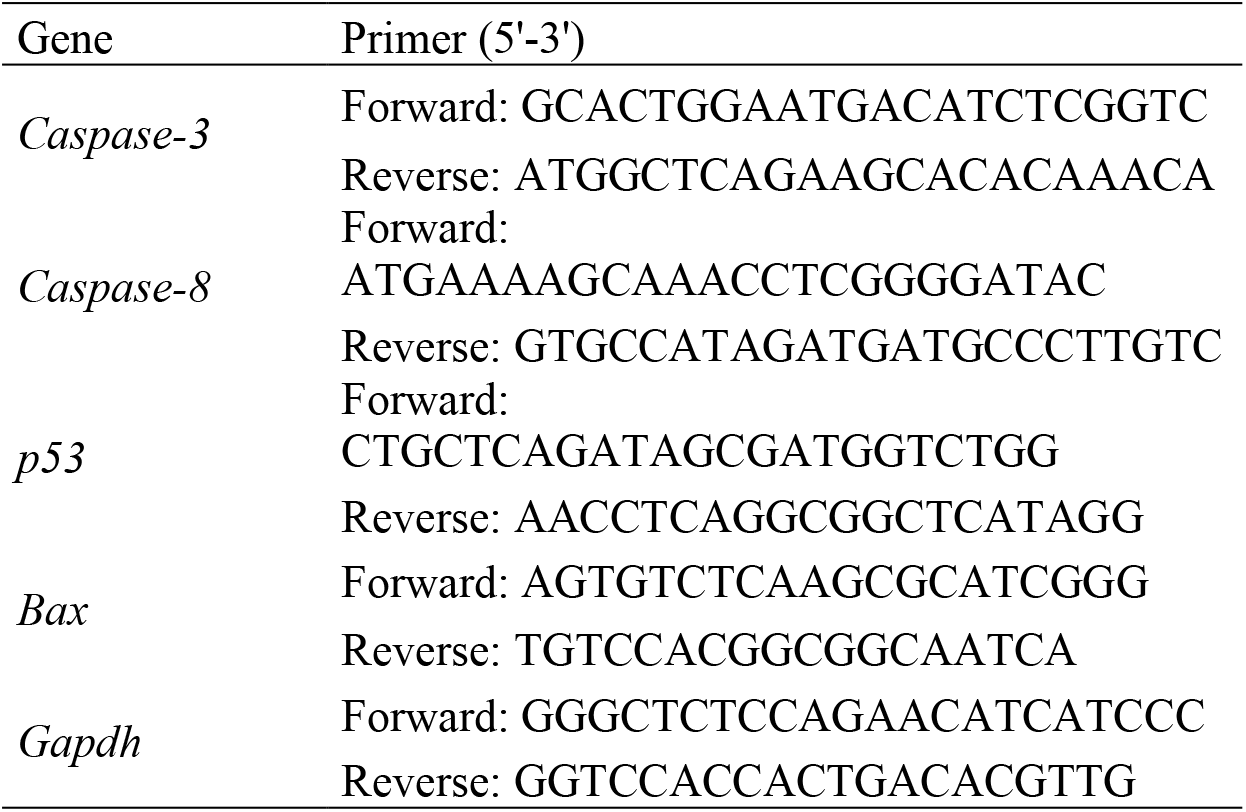
Primer sequences used in RT-qPCR

### TUNEL and LDH tests

The cells were plated into six-well tissue culture plates (Corning) at 10^6^ cells/well and treated with IC_50_ concentration of plant extracts. The next day apoptotic and necrotic cells were detected by flow cytometry (BD FACS Canto II) using APO-BrdU terminal deoxynucleotidyl transferase mediated dUTP nick end labeling (TUNEL) Assay Kit (Invitrogen) and lactate dehydrogenase (LDH) cytotoxicity assay kit (Pierce) according to the manufacturer’s protocol, respectively.

### Statistical Analysis

The statistical significance of the MTT test results was determined by One-way ANOVA test (SPSS, IBM, USA) and Bonferroni test. qRT-PCR results were evaluated using the Holm-Sidak method in Sigma Stat3.5 software. The statistical significance of the LDH test results was determined by student’s t-test. Values less than 0.05 (*) were considered to be statistically significant.

## Results

### *E. parviflorum* extracts reduce cell viability of HT-29 cell line

The human colon cancer cell line HT-29 and human noncancerous fibroblast cell line BJ were treated with 2 different *E. parviflorum* extracts which were extracted with water and ethanol. Each extract were given to the cells at various concentrations; 10, 30, 50, 100 μg/ml for water and 10, 100, 200, 500 μg/ml for ethanol. Viability of HT-29 cells was significantly reduced in a dose dependent manner 48 hr and 72 hr after administration of each extract (Figure 1, 2). Water and ethanol extracts reduced HT-29 cell viability up to 50% when applied at a dose of 43,6 μg/ml and 191.0 μg/ml, respectively (Figure 1A, 1C; Figure 2A, 2C; Figure 3). On the other hand, notably these concentrations did not significantly reduce the viability of noncancerous BJ fibroblast cells (Figure 1B; Figure 2B).

**Figure 1.**
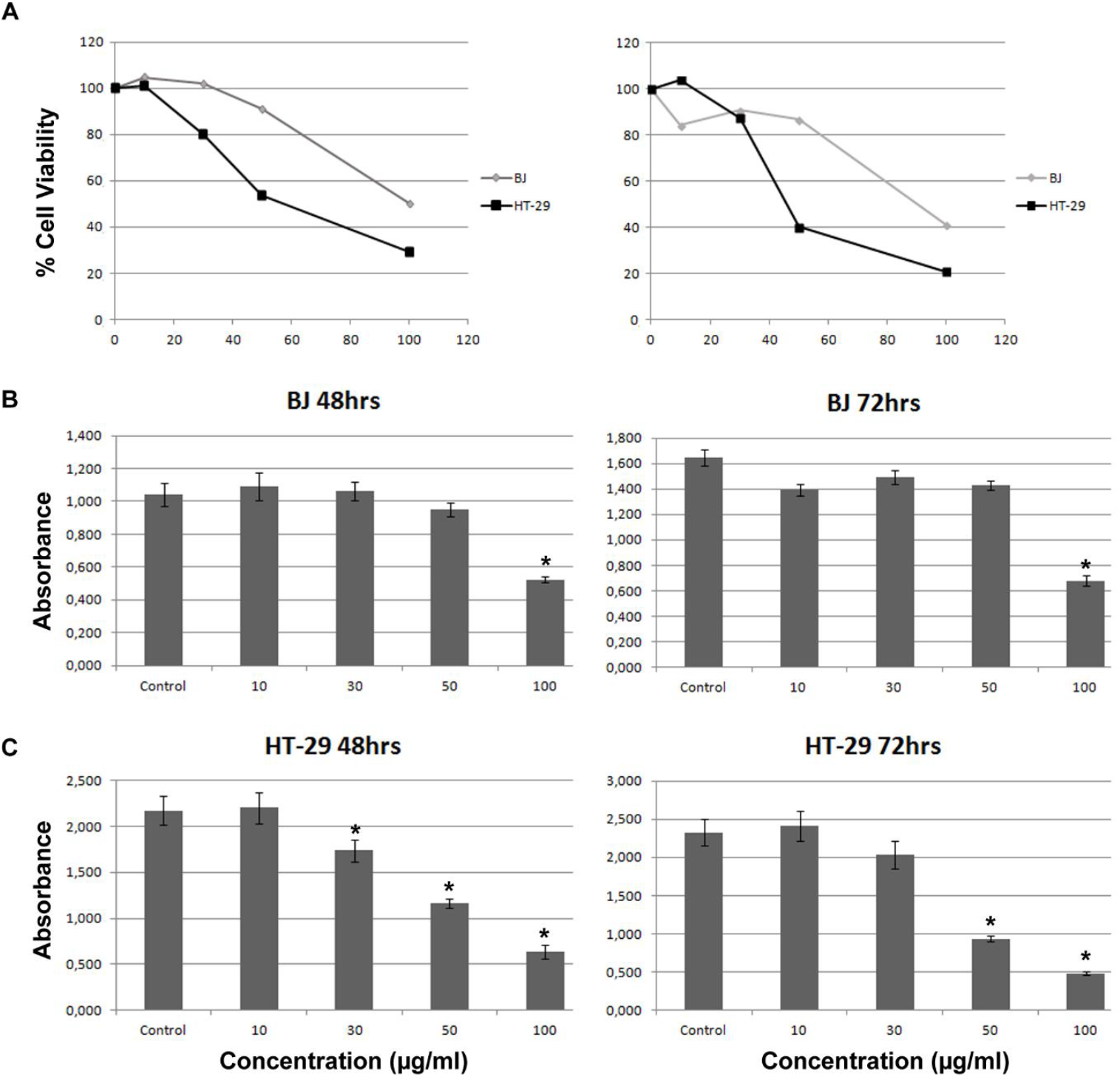
Cell viability percentage of HT-29 and BJ cells after 48 h (left panel) and 72 h (right panel) treatment with **aqueous extract of *E. parviflorum*** at different concentrations (μg/ml) **(A)**. Absorbance values from MTT assay of BJ **(B)** and HT-29 **(C)** cells after 48 h (left panel) and 72 h (right panel) treatment with aqueous extract of *E. parviflorum* at different concentrations (μg/ml). Values are expressed as mean ± S.D. (*n* = 3). Asterisk (*) indicates significant differences (*p* < 0.05).

**Figure 2.**
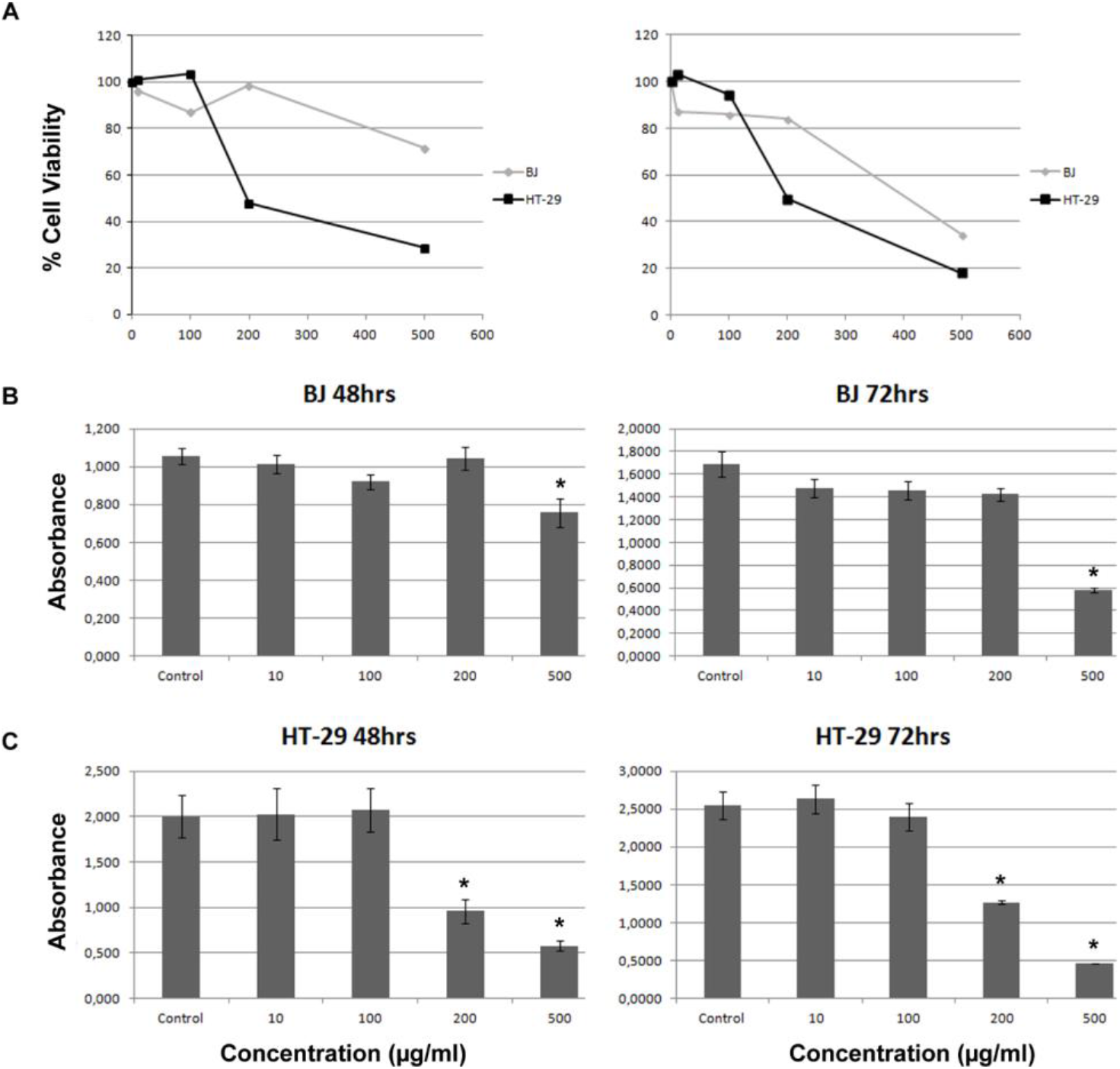
Cell viability percentage of HT-29 and BJ cells after 48 h (left panel) and 72 h (right panel) treatment with **ethanolic extract of *E. parviflorum*** at different concentrations (μg/ml) **(A)**. Absorbance values from MTT assay of BJ **(B)** and HT-29 **(C)** cells after 48 h (left panel) and 72 h (right panel) treatment with aqueous extract of *E. parviflorum* at different concentrations (μg/ml). Values are expressed as mean ± S.D. (n = 3). Asterisk (*) indicates significant differences (p < 0.05).

**Figure 3.**
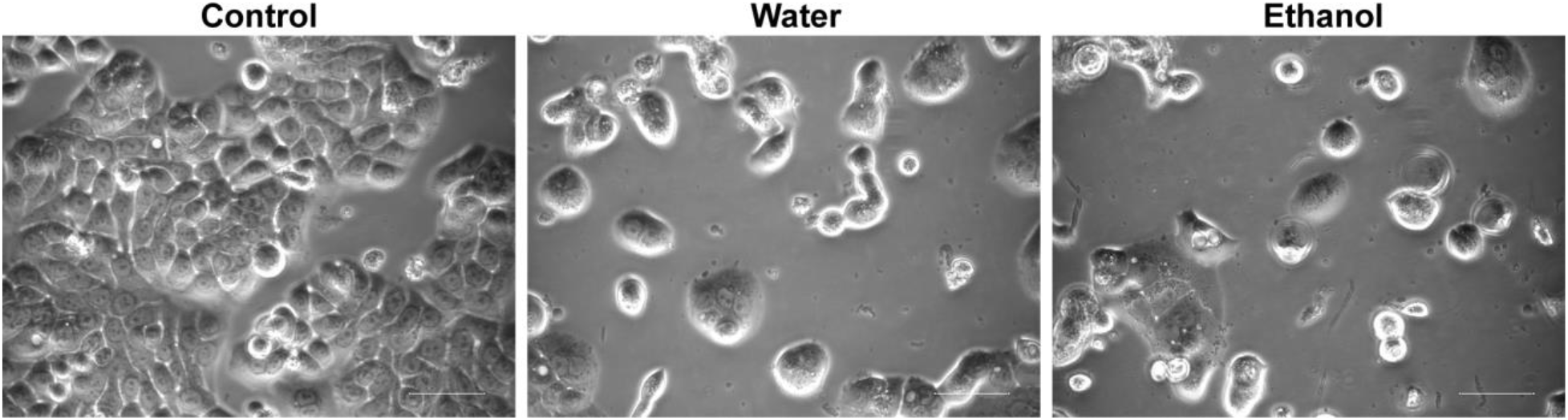
Reduced viability of HT-29 cells treated with aqueous and ethanolic extracts of *E. parviflorum*. Scale bars, 50μm.

### The effect of *E. parviflorum* extracts on gene expression profile

To further explore the effects of *Epilobium* extracts on HT-29 cells, the apoptotic gene expression profile of HT-29 and BJ cells were assessed 48 hr after administration of both extracts at their lethal concentration (LD_50_). For this purpose, mRNA expression levels of three different apoptotic genes (*Caspase-3*, *Caspase-8*, *Bax*), as well as the *p53* gene, were demonstrated through RT-qPCR (Figure 4). Results showed that both *E. parviflorum* extracts increased the expression level of apoptotic genes *Caspase-8* (1.72 ± 0.16-fold for ethanol, 1.82 ± 0.35-fold for water extracts) and *Bax* (1.74 ± 0.04-fold for ethanol, 2.45 ± 0.27-fold for water extracts) at different but significant degrees in HT-29 cells (Figure 4). However, both extracts significantly reduced the expression level of *p53* gene (0.57 ± 0.06-fold for ethanol, 0.41 ± 0.06-fold for water extracts) in HT-29 cells. However, the increase observed at apoptotic gene expression levels of HT-29 cells was not seen in BJ fibroblasts. Instead, their expression levels were reduced (0.34 ± 0.03, 0.52 ± 0.04, 0.37 ± 0.02-fold for *Caspase-8*, *Bax* and *Caspase-3*, respectively) (Figure 4). Aqueous extract, however, stimulated the expression of apoptotic genes (2.69 ± 0.34, 3.93 ± 0.17, 2.1 ± 0.14 for *Caspase-8*, *Bax* and *Caspase-3*, respectively) in BJ fibroblast cells which is consistent with 72 hr cell viability assays of aqueous extract on BJ fibroblast cells.

**Figure 4.**
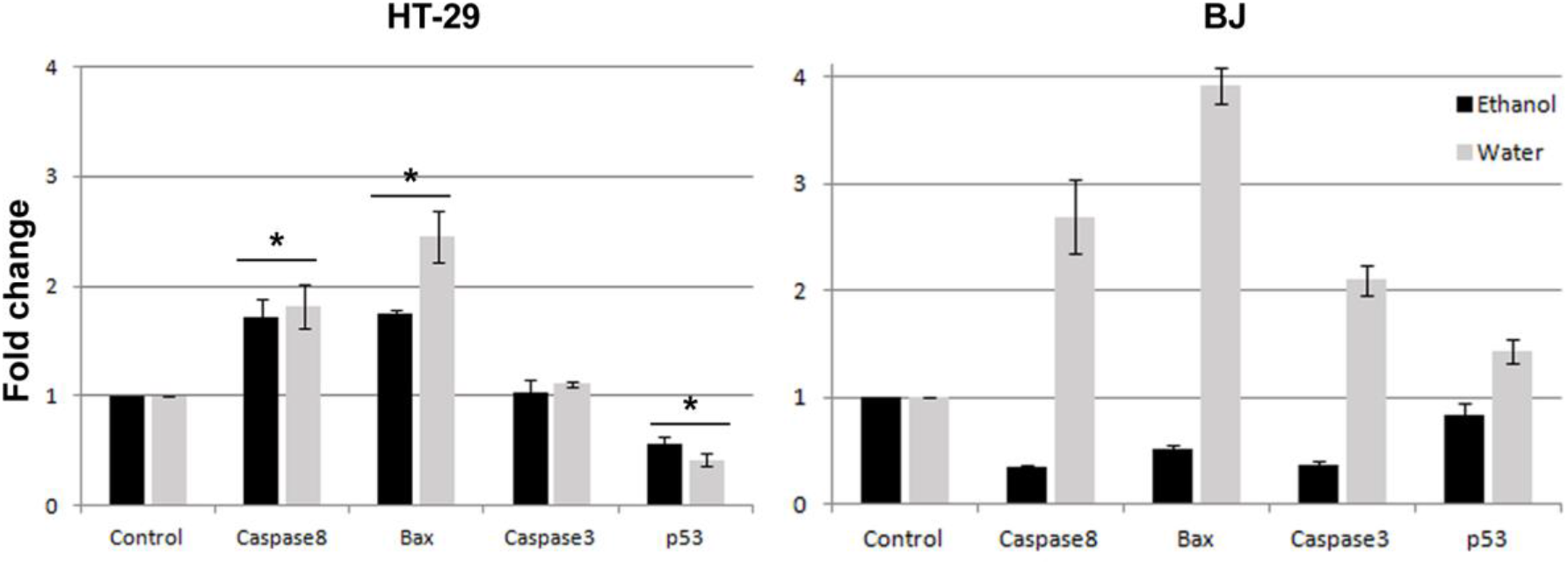
Apoptotic gene expression profile of HT-29 (left panel) and BJ (right panel) cells treated with ethanolic (black bars) and aqueous (grey bars) extracts of *E. parviflorum* for 48 hours. Data are represented as fold chance relative to untreated control cells. Each gene expression was normalized according to *Gapdh* expression. Values are expressed as mean ± S.D. (n = 3). Asterisk (*) indicates significant differences (p < 0.05).

### *E. parviflorum* extracts induce apoptotic cell death of HT-29 cells

In addition to profiling the apoptotic gene expression in HT-29 cells treated with *E. parviflorum* extracts, another morphological sign of apoptosis (nuclear fragmentation) in these cells was demonstrated by performing TUNEL assay. After labeling the 3’-OH edges of fragmented nuclear DNA through a TUNEL assay kit, flow cytometry analysis showed that 6.9% and 13.3% of HT-29 cells were fated to apoptosis when treated with water and ethanol extracts respectively (Figure 5A). Nuclear fragmentation in these cells was also demonstrated through immunostaining under a fluorescence microscope (Figure 5B). Moreover, LDH assay showed that both *E. parviflorum* extracts slightly but significantly stimulated the necrosis in HT-29 cells compared to untreated control cells (Figure 6).

**Figure 5.**
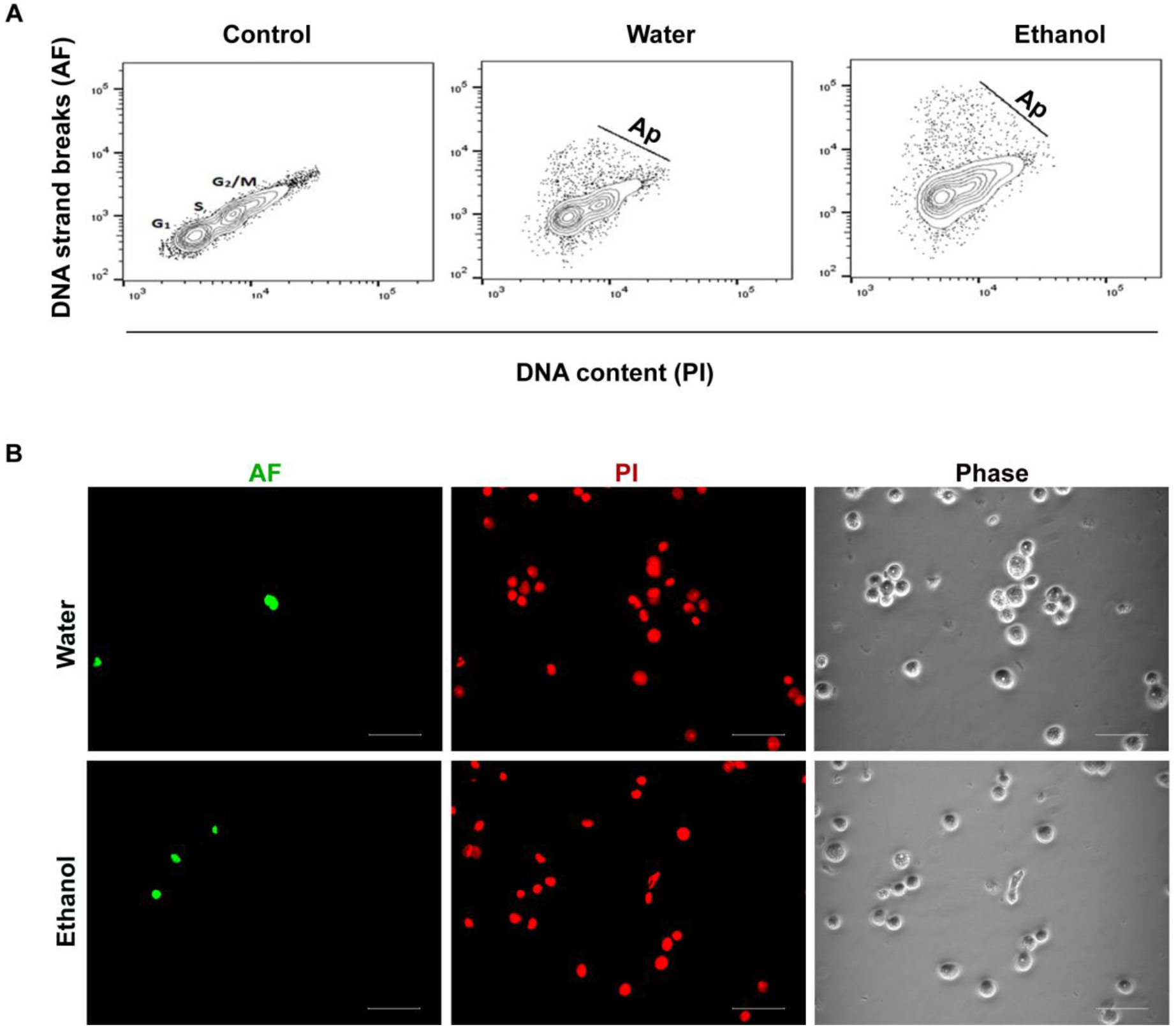
Demonstration of apoptotic HT-29 cells treated with *E. parviflorum* extracts through FACS **(A)** and fluorescence microscope **(B)**. Cells were analyzed after 24 h extract treatment. DNA strand breaks in apoptotic cells were labeled with Alexa Fluor 488 (AF; green). Cell nuclei of all cells were labeled with propidium iodide (PI; red). Scale bars, 50μm.

**Figure 6.**
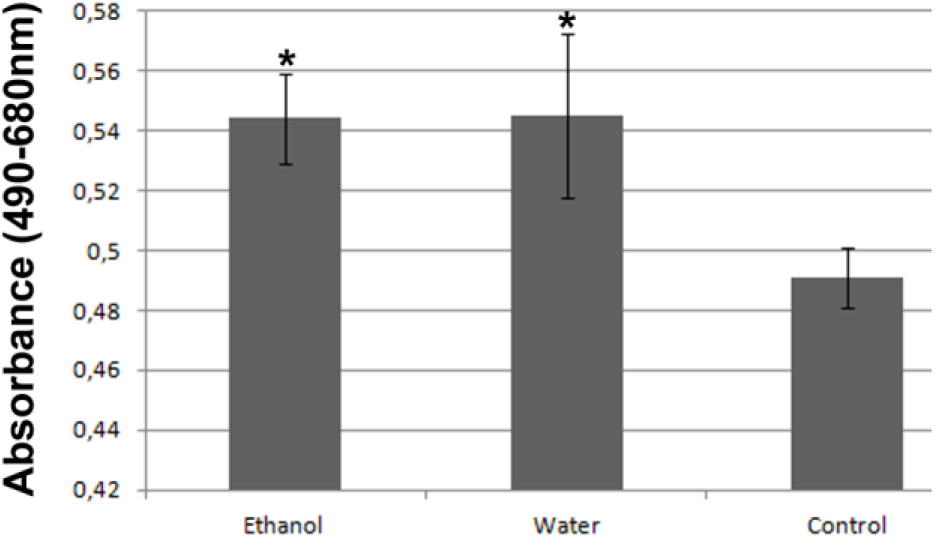
Necrosis of HT-29 cells treated with *E. parviflorum* extracts. Cells were analyzed after 24 h extract treatment. Values are expressed as mean ± S.D. (n = 3). Asterisk (*) indicates significant differences (p < 0.05).

In consistent with being widely used in folk medicine, *Epilobium* spp. have been demonstrated to have various therapeutic effects such as analgesic (Pourmorad et al. 2007; Tita et al. 2001), antiandrogenic (Allkanjari and Vitalone 2015), anti-inflammatory (Hevesi et al. 2009; Vogl et al. 2013), antimicrobial (Battinelli et al. 2001; Jones et al. 2000), antioxidant (Stajner et al. 2007; Steenkamp et al. 2006), and antitumor activity (Vitalone et al. 2003a; Voynova et al. 1991) in various *in vitro* as well as *in vivo* animal studies. Antiproliferative effect of *Epilobium* extracts on healthy (PZ-HPV-7) and cancerous prostate cells lines (LNCaP) was also reported by different groups and this effect is mainly associated with its ellagitannins; Oenothein A and B which inhibit aromatase and 5-α-reductase enzymes involved in etiology of benign prostatic hyperplasia (Ducrey et al. 1997; Lesuisse et al. 1996; Piwowarski et al. 2017; Yoshida et al. 2018). However, demonstration of anti-tumorigenic and antiproliferative effect of *Epilobium* extracts on leucosis (P388) (Voynova et al. 1991) and astrocytoma (1321N1) (Vitalone et al. 2003b) cell lines respectively indicates that this effect is not specific for prostate cells. Which biological compounds of *Epilobium* extracts are responsible for this antiproliferative effect is not known yet, even though there are studies demonstrating the antiproliferative effect of ellagitannins on different cancer types (Ismail et al. 2016; Wang et al. 1999).

## Discussion

Our results from the present study demonstrated that different *E. parviflorum* extracts, which are prepared by using water and ethanol solvents, reduced the viability of HT-29 cells significantly (Figure 1; Figure 2; Figure 3). The potency of plant extracts depends on the extraction method. The most potent extraction solvent also depends on the plant species to be studied as well as the active molecule to be isolated. In the present study the aqueous extract (43,6 μg/ml) of *E. parviflorum* is more potent than the ethanolic extract (191.0 μg/ml) on HT-29 cells (Figure 1; Figure 2). This may come from the different degree of polarity of these solvents. Ethanol is less polar than water and more hydrophilic components are known to be retained in aqueous extracts compared to ethanolic extracts (Abarca-Vargas et al. 2016; Rauf et al. 2018). On the other hand, it should also be noted that ethanol is a better solvent for organic compounds than the water is. Another reason why water extract was more potent may be due the fact that we re-suspended both extracts in water which may not dissolve the non-polar components extracted with ethanol. It is also notable that these concentrations reducing the viability of HT-29 cells up to 50% did not have much effect on healthy fibroblast cells. RT-qPCR results showed that both extracts increased the expression of apoptotic genes *Caspase-8* and *Bax* (Figure 4). The tunel assay, in which 3’-OH ends of fragmented DNAs are fluorescently labeled, also verified cell death through apoptosis (Figure 5). Apoptosis can be initiated by intrinsic and extrinsic pathways. Extrinsic pathways activate Caspase-8 whereas intrinsic pathways activate Caspase-9; both of which are called initiator factors, and can separately induce executioner factors to initiate apoptosis (McIlwain et al. 2013). Bax is also a regulator of intrinsic pathways of apoptosis and induces mitochondrial dysfunction by mitochondrial membrane permeabilization (Liu et al. 2016). Permeabilization of mitochondrial membrane causes the release of cytochrome c which in turn stimulates apoptosis through Caspase-9 (Liu et al. 2016). Our RT-qPCR results indicate that *E. parviflorum* extracts may induce apoptosis both through intrinsic and extrinsic pathways. Consistent with our results, Stolarczyk et al. (2013) also demonstrated the apoptotic effect of different *Epilobium* extracts on human hormone dependent prostate cancer cells through intrinsic pathways (Stolarczyk et al. 2013). In addition to stimulation of apoptosis, we also found that HT-29 cells treated with *E. parviflorum* extracts also underwent necrosis (Figure 6). The observed necrosis could be secondary necrosis due to the toxic content of apoptotic cells which are not cleaned out by macrophages in *in vitro* culture (Silva 2010). On the other hand, there are also reports showing that the same compounds can even cause apoptosis and necrosis simultaneously (Arakawa et al. 2015). We also showed that expression of *p53* gene is downregulated in HT-29 cells treated with *E. parviflorum* extracts. Wild type p53 is a tumor suppressor protein and *p53* gene is mutated in many cancer cell types including HT-29 cells (Freed-Pastor and Prives 2012; Levine and Oren 2009). In HT-29 cells, *p53* gene has a R273H mutation (Rodrigues et al. 1990). There are many reports demonstrating that R273H substitution in *p53* is a gain of function mutation and providing oncogenic functions to the cells to drive invasion and metastasis (Lang et al. 2004; Muller et al. 2013; Olive et al. 2004; Sigal and Rotter 2000). Based on the literature, reduction of mutant *p53* expression, as we saw in our study, may be useful to reduce the oncogenic properties of HT-29 cells as well.

## Conclusion

Our results confirmed that *E. parviflorum* extracts can eliminate proliferation of human colorectal carcinoma cells *in vitro*. The findings we obtained indicate that aqueous and ethanolic extracts of *E. parviflorum* have the potential to become therapeutic anti-carcinogenic agent. However, to support the hypothesis that *E. parviflorum* extracts might play a role in the colorectal cancer prevention, some randomized controlled trials as well as large prospective cohort studies are needed.

## Acknowledgements

The authors thank Dr. Trevor A. Epp for his helpful comments on the manuscript.

## Author contribution statement

MAA and EA conceived the study; TS, EA, NE and MAA conducted the experiments. EA and MAA wrote the manuscript, and all authors read, edited and approved the manuscript.

